# RepeatFiller newly identifies megabases of aligning repetitive sequences and improves annotations of conserved non-exonic elements

**DOI:** 10.1101/696922

**Authors:** Ekaterina Osipova, Nikolai Hecker, Michael Hiller

## Abstract

Transposons and other repetitive sequences make up a large part of complex genomes. Repetitive sequences can be co-opted into a variety of functions and thus provide a source for evolutionary novelty. However, comprehensively detecting ancestral repeats that align between species is difficult since considering all repeat-overlapping seeds in alignment methods that rely on the seed-and-extend heuristic results in prohibitively high runtimes. Here, we show that ignoring repeat-overlapping alignment seeds when aligning entire genomes misses numerous alignments between repetitive elements. We present a tool – RepeatFiller – that improves genome alignments by incorporating previously-undetected local alignments between repetitive sequences. By applying RepeatFiller to genome alignments between human and 20 other representative mammals, we uncover between 22 and 84 megabases of previously-undetected alignments that mostly overlap transposable elements. We further show that the increased alignment coverage improves the annotation of conserved non-exonic elements, both by discovering numerous novel transposon-derived elements that evolve under constraint and by removing thousands of elements that are not under constraint in placental mammals. In conclusion, RepeatFiller contributes to comprehensively aligning repetitive genomic regions, which facilitates studying transposon co-option and genome evolution.

**Source code:** https://github.com/hillerlab/GenomeAlignmentTools

## Introduction

A substantial portion of vertebrate genomes consist of transposons and other repetitive sequences [1, 2]. While most repeats are estimated to evolve neutrally [3], transposons are important substrates for evolutionary tinkering [4, 5]. For example, transposon-derived sequences contribute to the transcriptome by providing alternatively spliced exons [6, 7]. By contributing transcription factor binding sites, promoters, and distal regulatory elements, co-opted transposons are involved in rewiring of regulatory networks and drive regulatory innovation [7-15]. Importantly, a sizeable portion of evolutionarily constrained regions arose from ancestral transposon sequences [16, 17]. Studying how ancestral transposons and other repeats were co-opted into functional roles requires whole genome alignments that comprehensively align orthologous repeats.

The nature of repetitive sequences such as transposons, however, leads to many paralogous alignments, which pose a challenge for comprehensively aligning orthologous repeats between vertebrate genomes. Most methods for aligning entire genomes use a seed-and-extend heuristic, originally implemented in BLAST [18], to find local alignments between the sequences of two genomes. The seeding step of this heuristic detects short words or patterns (called seeds) that match between the sequences of the two genomes. This can be computed very efficiently. Seed detection is then followed by a computationally more expensive alignment extension step that considers ungapped and gapped local alignments. Given that repetitive sequences provide numerous seed matches to paralogous repeat copies in a whole genome comparison, it is computationally infeasible to start a local alignment from seeds located in repetitive sequences. Therefore, seeds that overlap repetitive regions are not used to start a local alignment phase, either by masking repetitive regions before aligning genomes [19-22] or by dynamically adapting seeding parameters by the observed seed frequencies [23]. Consequently, alignments between highly-identical repeats are only found during the extension phase, initiated from seeds outside the repeat boundaries. This can be problematic if the regions flanking a repeat have been diverged to an extent that no seed in the vicinity of the repeat can be found.

Here, we investigated to which extent aligning repetitive sequences are missed in whole genome alignments. We show that ignoring repeat-overlapping seeds misses between 22 and 84 Mb of mostly repetitive elements that actually align between mammals and we provide a tool, called RepeatFiller, to incorporate such repeat-overlapping alignments into genome alignments. We further show that a subset of aligning sequences detected by RepeatFiller evolve under evolutionary constraint, which uncovers previously-unknown conserved non-exonic elements and thus improves the annotation of constrained elements.

## Results

### RepeatFiller incorporates several megabases of aligning repetitive sequences to genome alignments

To investigate how many aligning repetitive elements have been missed in alignments between mammalian genomes, we adopted a previously-developed approach that was initially devised to detect novel local alignments between a pair of distantly-related species [24, 25]. The original approach focused on unaligning regions that are flanked by aligning blocks in co-linear alignment chains [26], which are detected in the first all-vs-all genome alignment step. In a second step, this original approach used lastz [21] with highly-sensitive seeding and (un)gapped extension parameters to align the previously-unaligning regions again. This second round of highly-sensitive local alignment can uncover novel alignments that are co-linear with already-detected alignment blocks. Here, we adopted this approach by introducing two key changes. First, we increased alignment parameter sensitivity only slightly, but unmasked the unaligning region. This implies that all seeds, including repeat-overlapping seeds, will be considered (Figure 1). By restricting the size of the unaligning regions to smaller regions of at most 20 kb, we reason that novel local alignments detected with a similar sensitivity level likely constitute orthologous alignments. Second, while the previous approach computed all alignment chains again from scratch using previously-detected and novel local alignments, our new approach directly adds novel alignments to existing alignment chains, thus removing the need for a chain re-computing step. This approach is called RepeatFiller and is available at https://github.com/hillerlab/GenomeAlignmentTools.

**Figure 1:**
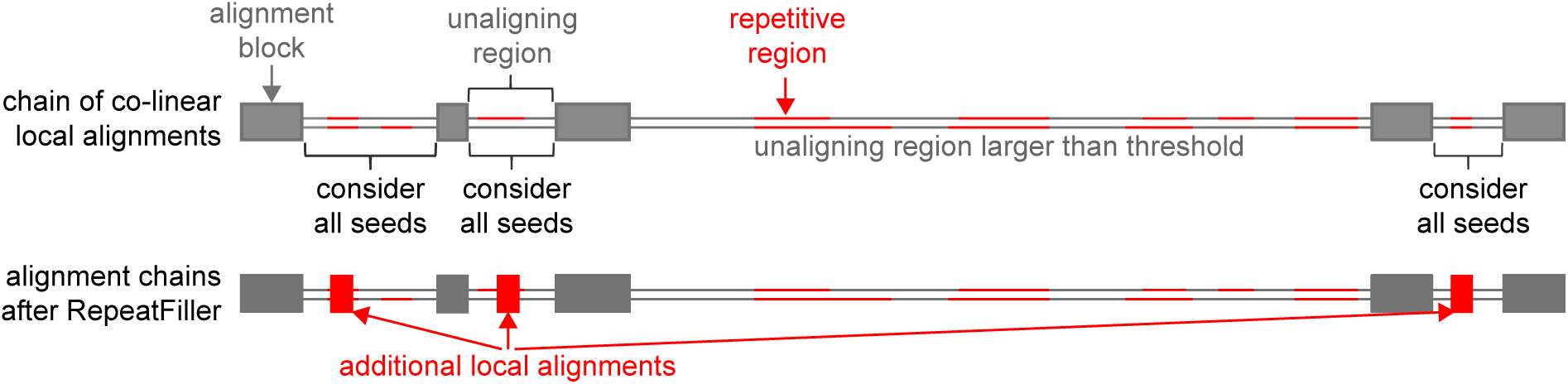
Missed repeat-overlapping alignments and concept of RepeatFiller. Illustration of RepeatFiller. Focusing on unaligning regions in a reference and query genome that are flanked by up-and downstream aligning blocks, RepeatFiller performs a second round of local alignment considering also repeat-overlapping seeds. Newly found local alignments (red boxes) are inserted into the context of other aligning blocks (grey boxes). Unaligning regions that are larger than a user-defined threshold are not considered as the chance of aligning non-orthologous repeats is increased.

To investigate how many aligning repetitive elements can be added by RepeatFiller, we built alignment chains between the human (hg38) genome assembly and the genomes of 20 other mammals that represent the major mammalian clades (Figure 2, Supplementary Table 1). We found that RepeatFiller adds between 22.4 Mb (Rhesus macaque) and 83.7 Mb (rabbit) of aligning sequence, which represents between 0.7 – 2.6% of the human genome (Figure 2, Supplementary Table 1). By overlapping the new alignments with repetitive elements annotated in the human genome, we found that the vast majority of newly-aligned sequences overlap repeats, in particular transposable elements (Figure 2, Supplementary Table 1). The runtime of the RepeatFiller step is between 14.7 and 43.4 CPU hours (Supplementary Table 1), and thus adds little the runtime of the initial genome-wide all-vs-all pairwise alignment step that is typically around ∼1000 CPU hours. Together, this shows that a considerable portion of aligning transposon sequences are missed when repeat-overlapping seeds are ignored and that RepeatFiller can detect such alignments with little extra computational runtime.

**Figure 2:**
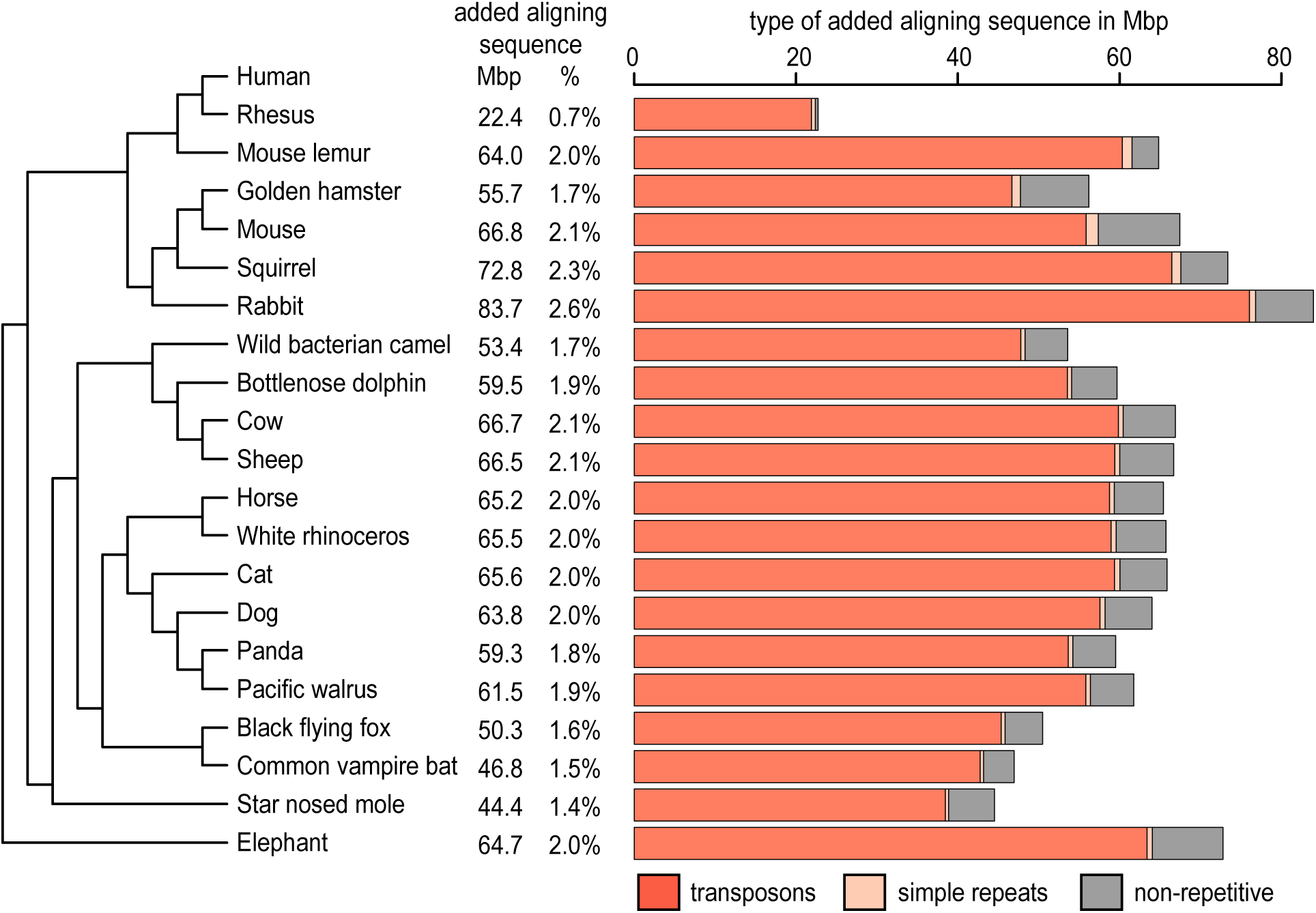
RepeatFiller adds several megabases of aligning transposable elements to existing genome alignments. Phylogenetic tree of human and 20 non-human mammals whose genomes we aligned to the human genome. The amount of newly alignments detected by RepeatFiller is shown in megabases and in percent relative to the human genome. Bar charts provide a breakdown of newly-added aligning sequences into overlap with transposons, simple repeats and non-repetitive sequence.

### RepeatFiller application uncovers thousands of novel repeat-derived conserved non-exonic elements

Next, we investigated whether some of the newly-aligning sequences show evidence of evolutionary constraint, which indicates purifying selection and a biological function. To this end, we used the pairwise alignments, generated either with or without RepeatFiller, to build two human-referenced multiple genome alignments of 21 mammals with Multiz [27]. Then, we used PhastCons [28] to identify constrained elements. We found that the majority (98%) of the 164 Mb in the human genome that are classified as constrained in the multiple alignment without RepeatFiller were also classified as constrained in the RepeatFiller-subjected alignment.

Dividing the conserved regions detected in the alignment without RepeatFiller into exonic and non-exonic regions, we found that 99.8% of the exonic and 97.4% of the non-exonic regions are also classified as constrained in the RepeatFiller-subjected alignment. Since conserved exonic regions are virtually identical, likely because they rarely overlap repeats, we focused our comparison on the conserved non-exonic elements (CNEs), which often overlap *cis*-regulatory elements [29-31]. This comparison first showed that 3.46 Mb of the human genome were newly classified as conserved non-exonic in the RepeatFiller-subjected alignment, representing 2.9% of all conserved non-exonic bases detected in this alignment. Requiring a minimum size of 30 bp, application RepeatFiller led to the identification of 30167 novel CNEs that are listed in Supplementary Table 2.

Two striking examples of newly-identified CNEs are shown in Figures 3 and 4. Figure 3 shows the genomic region overlapping *MEIS3*, a homeobox transcription factor gene that synergizes with Hox genes and is required for hindbrain development and survival of pancreatic beta-cells [32-34]. By revealing novel alignments to many non-human mammals, RepeatFiller identifies several novel repeat-overlapping CNEs in introns of *MEIS3* (Figure 3). Figure 4 shows the genomic region around *AUTS2*, a transcriptional regulator required for neurodevelopment that is associated with human neurological disorders such as autism [35, 36]. Applying RepeatFiller revealed several novel CNEs upstream of *AUTS2*. For some of these CNEs, RepeatFiller incorporated a well-aligning sequence of 19 mammals, which then permitted the identification of evolutionary constraint. Overall, applying RepeatFiller led the identification of more than 30000 CNEs that were not detected before.

**Figure 3:**
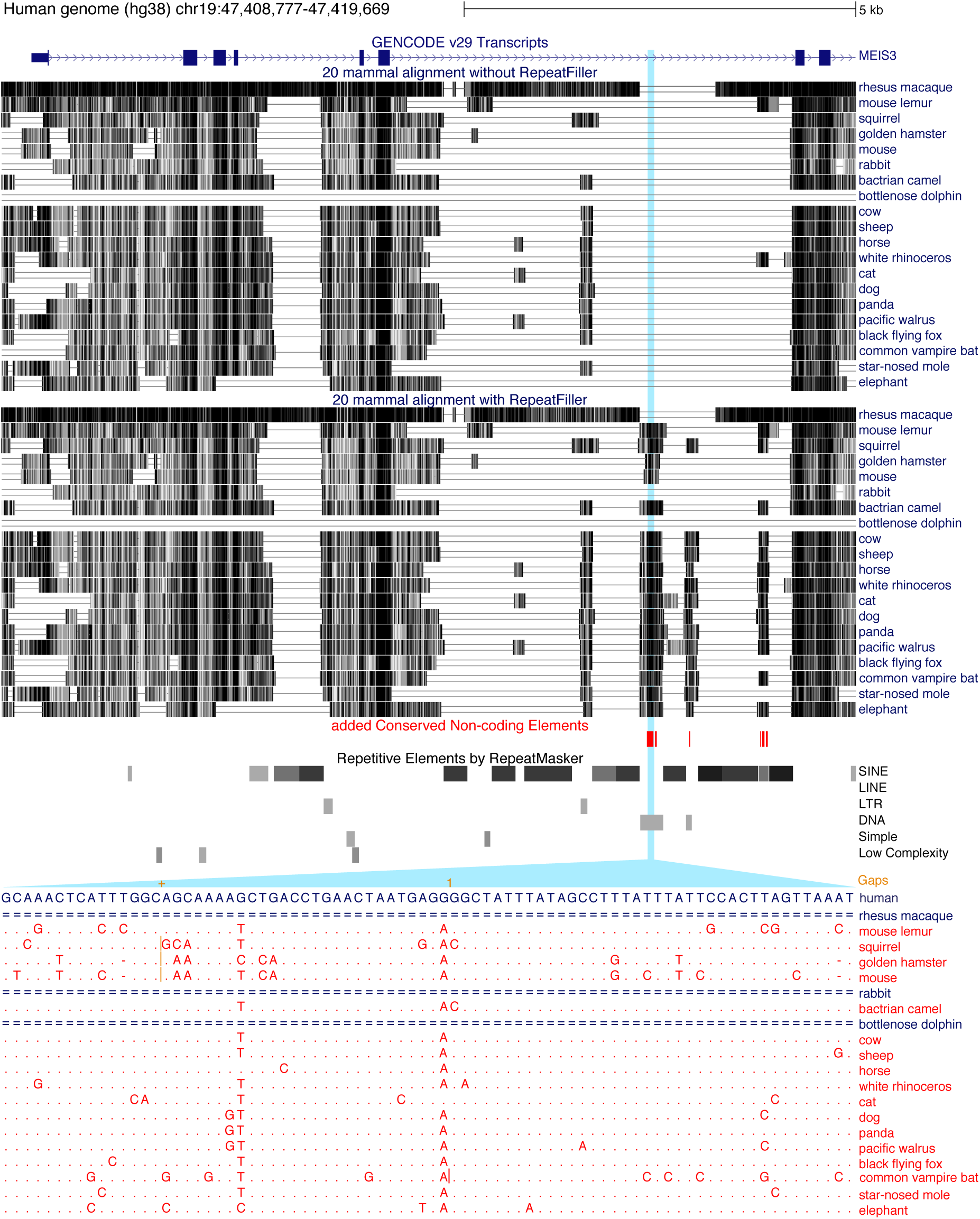
Examples of newly-identified CNEs near MEIS3. UCSC genome browser [38] screenshot shows an ∼11 kb genomic region overlapping the gene MEIS3, a homeobox transcription factor that is required for hindbrain development. Visualization of the two multiple genome alignments (without RepeatFiller at the top, with RepeatFiller below; boxes representing align regions with darker colors indicating a higher alignment identity) shows that RepeatFiller adds several aligning sequences, some of which evolve under evolutionary constraint and thus are CNEs (red boxes) only detected in the RepeatFiller-subjected alignment. The RepeatMasker annotation shows that these newly-identified CNEs overlap transposons. The zoom-in shows the 21-mammal alignment of one of the newly-identified CNEs, which overlaps a DNA transposon. While this genomic region did not align to any mammal before applying RepeatFiller, our tool identified a well-aligning sequence for 17 non-human mammals (red font). A dot represents a base that is identical to the human base, insertions are marked by vertical orange lines, and unaligning regions are showed as double lines.

**Figure 4:**
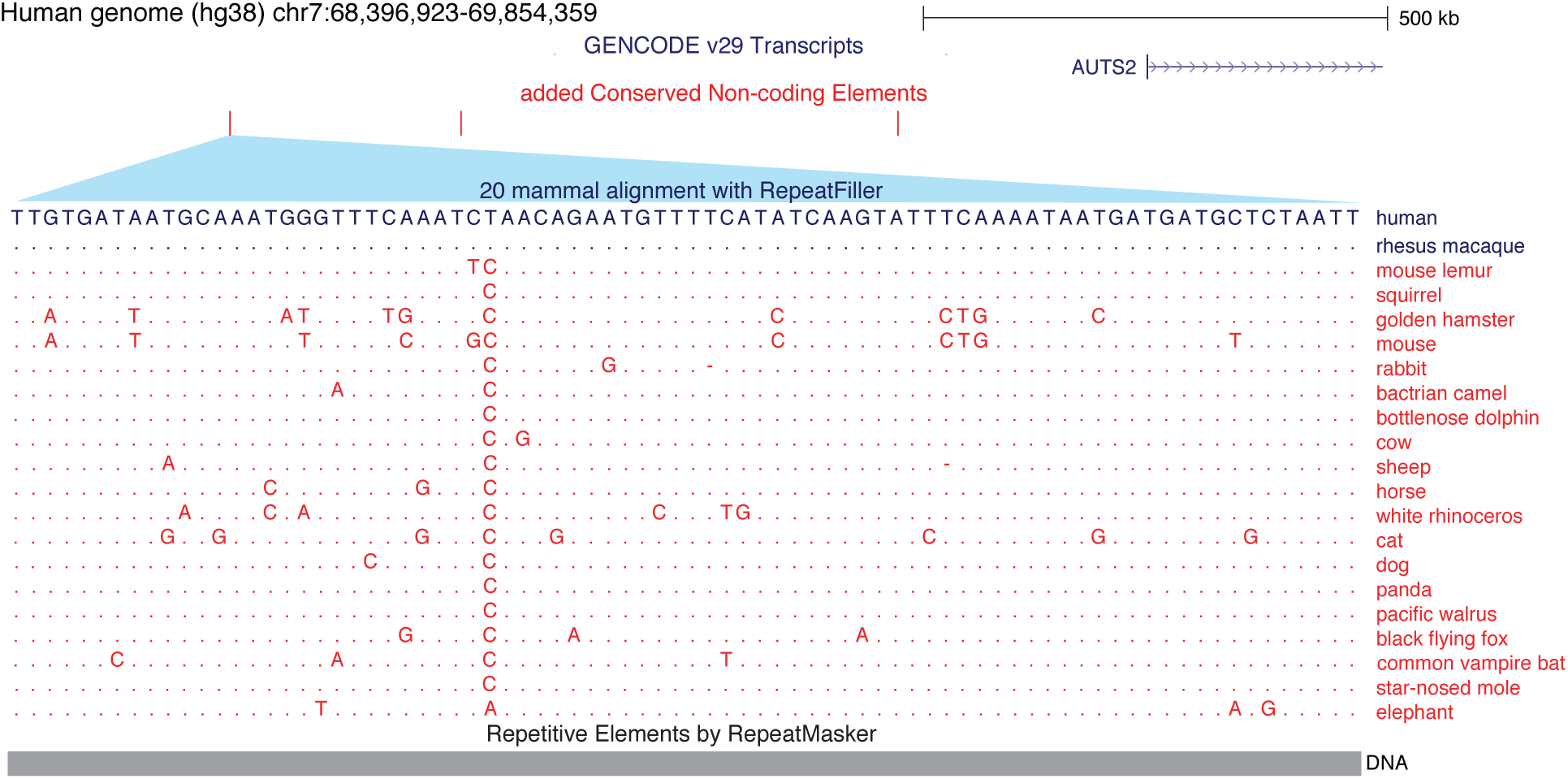
Examples of newly-identified CNEs upstream of *AUTS2*. UCSC genome browser screenshot shows a ∼1.5 Mb genomic region around *AUTS2*, a transcriptional regulator required for neurodevelopment. CNEs only detected in the RepeatFiller-subjected multiple alignment are marked as red tick marks. The zoom-in shows the 21-mammal alignment of one of the newly-identified CNEs. While only the rhesus macaque sequence aligned to human before applying RepeatFiller, our tool identifies a well-aligning sequence for all 19 other mammals (red font). A dot represents a base that is identical to the human base. The RepeatMasker annotation (bottom) shows that this newly-identified CNE overlaps a DNA transposon.

### RepeatFiller improves annotations of Conserved Non-exonic Elements

Interestingly, the comparison of conserved non-exonic bases detected by PhastCons also revealed 3.08 Mb of the human genome that were classified as conserved non-exonic only in the multiple alignment without RepeatFiller, but not in the RepeatFiller-subjected alignment. These 3.08 Mb represent 2.6% of all conserved non-exonic bases detected in the alignment without RepeatFiller. The 29334 CNEs with a size ≥30 bp are listed in Supplementary Table 3. To investigate the reasons underlying these ‘lost’ CNEs, we first sought to confirm that the RepeatFiller-subjected alignment had an increased species coverage in these regions. Indeed, we found that RepeatFiller added on average 3.9 (median 3) aligning species to these lost CNEs. Inspecting many of these CNEs showed that the newly added sequences are similar to the already-aligned sequences; however, they exhibit more substitutions. These substitutions increase the overall sequence divergence across mammals, which likely explains why the same region was not classified as constrained anymore, despite having a higher coverage of aligning species. Figure 5 A and B shows two examples of such genomic regions that are not classified as constrained after adding additional alignments with RepeatFiller.

**Figure 5:**
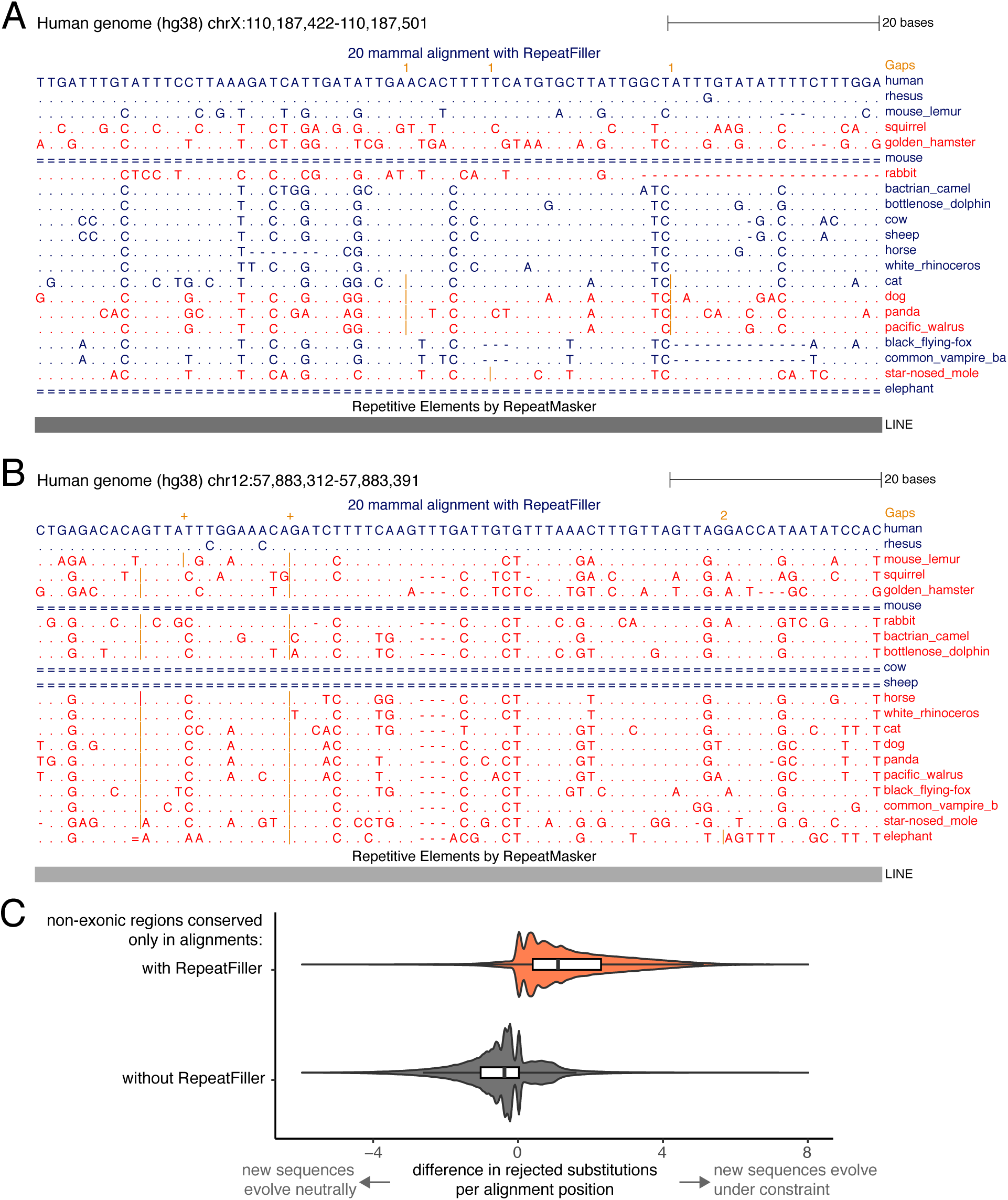
Additional alignments found with RepeatFiller reveal absence of conservation in the genomic regions that were erroneously classified as conserved before. (A, B) UCSC genome browser screenshots showing two examples of genomic regions that were only classified as constrained in a multiple genome alignment generated without applying RepeatFiller. Dots in these alignments represent bases that are identical to the human base, insertions are marked by vertical orange lines, and unaligning regions are showed as double lines. The alignments show that the sequences of species added by RepeatFiller (red font) exhibit a number of substitutions. This explains why these regions were not classified as constrained anymore, despite adding more aligning sequences. Please note that in (B) only the sequence of the rhesus macaque was aligned before applying RepeatFiller. Sequences in both (A) and (B) overlap LINE transposons. (C) Difference in evolutionary constraint in non-exonic alignment columns that are only classified as constrained in either alignment. For each alignment position, we used GERP++ to compute the estimated number of substitutions rejected by purifying selection (RS). The difference in RS between alignments with and without RepeatFiller is visualized as a violin plot overlaid with a white box plot. This shows that almost all non-exonic bases that were only detected as constrained in the alignment with RepeatFiller (orange background) have a positive RS difference, indicating that the newly-aligning sequences added by RepeatFiller largely evolve under evolutionary constraint. In contrast, non-exonic bases only detected as constrained in the alignment without RepeatFiller (grey background) often have slightly negative RS differences, indicating that many of the newly-added sequences do not evolve under constraint. The two distributions are significantly different (P<e^−16^, two-sided Wilcoxon rank sum test).

To confirm that the newly-added sequences increase the overall sequence divergence, we applied GERP++ [37] to both multiple alignments (Supplementary Figure 1A). For each alignment column, GERP++ estimates the number of substitutions that were rejected by purifying selection (RS = rejected substitutions) by subtracting the number of observed substitutions from the number of substitutions expected under neutrality. Since GERP++ computes the number of substitutions expected under neutrality from a phylogenetic tree that is pruned to the aligning species (Supplementary Figure 1B), we can directly compare RS between alignment columns that were only classified as constrained in either alignment to estimate whether the RepeatFiller-added sequences evolve slower than expected under neutrality. Specifically, for each alignment column, we computed the difference in RS before and after adding new alignments with RepeatFiller, as illustrated in Supplementary Figure 1B.

We found that the alignment columns, where constraint was only detected in the alignment without RepeatFiller, mostly exhibit slightly negative RS differences (Figure 5C, grey background), which suggests that many positions in the RepeatFiller-added sequences do not evolve under strong constraint. Hence, the extent of constraint in the more limited set of aligning sequences was likely overestimated, providing an explanation of why these genomic regions were not classified anymore as constrained across placental mammals. It should be noted that these regions may still be under constraint in particular lineages. In contrast, most alignment columns, where constraint was only detected after applying RepeatFiller, exhibit a positive RS difference (Figure 5C, orange background), which suggests that the newly-added sequences evolve under constraint. Overall, by uncovering previously-unknown alignments, RepeatFiller application led to an improved CNE annotation.

## Discussion

While transposon-derived sequences can be co-opted into a multitude of biological roles and can evolve under evolutionary constraint, comprehensively detecting alignments between ancestral transposons and other repeats is not straightforward. The main reason is that considering all repeat-overlapping alignment seeds during the initial whole genome alignment step is computationally not feasible. However, it is feasible to consider all seeds when aligning local regions that are bounded by colinear aligning blocks. We provide a tool RepeatFiller that implements this idea and incorporates newly-detected repeat-overlapping alignments into pairwise alignment chains. We tested the tool on alignments between human and 20 representative mammals and showed that with little additional computational runtime RepeatFiller uncovers between 22 and 84 Mb of previously-undetected alignments that mostly originate from transposable elements.

We further show that RepeatFiller application enables a refined and more complete CNE annotation by two means. First, applying RepeatFiller led the identification of thousands of CNEs whose aligning sequences were not detected before. This includes highly-conserved transposon-derived CNEs that are located near important developmental genes. Second, the sequences added by RepeatFiller may not evolve slower than expected under neutral evolution. In this case, providing a more complete set of aligning sequences led to the removal of thousands of putatively-spurious CNEs that overall do not evolve under strong constraint across placental mammals, though the possibility of lineage-specific constraint remains.

Taken together, RepeatFiller implements an efficient way to improve the completeness of aligning repetitive regions in whole genome alignments, which helps annotating conserved non-exonic elements and studying transposon co-option and genome evolution.

## Materials and Methods

### Generating pairwise genome alignments

We used the human hg38 genome assembly as the reference genome. To compute pairwise genome alignments, we used lastz version 1.04.00 [21] and the chain/net pipeline [26] with default parameters (chainMinScore 1000, chainLinearGap loose). We used the lastz alignment parameters K = 2400, L = 3000, Y = 9400, H = 2000 and the lastz default scoring matrix. All species names and their assemblies are listed in Supplementary Table 1.

### RepeatFiller

The input of RepeatFiller is a file containing co-linear chains of local alignment blocks. This file must be in the UCSC chain format as defined here https://genome.ucsc.edu/goldenPath/help/chain.html. The output is a file that contains the same chains plus the newly-added local alignment blocks. By default, RepeatFiller only considers unaligned regions in both the reference and query genome that are at least 30 bp and at most 20000 bp long. We considered all chains with the score greater than 25000. For each unaligning region that fulfills the size thresholds, RepeatFiller uses lastz with the same parameters as above but with a slightly more sensitive ungapped alignment threshold (K=2000). All repeat-masking (lower case letters) was removed before providing the local sequences to lastz. Since lastz may find multiple additional local alignments in this second step, we used axtChain [26] to obtain a ‘mini chain’ of local alignments for this unaligning region. RepeatFiller then inserts the aligning blocks of a newly-detected mini chain at the respective position in the original chain if the score of the mini chain is at least 5000. All default parameters for the size of unaligning regions, minimum chain scores and local alignment parameters can be changed by the user via parameters. Finally, RepeatFiller recomputes the score of the entire chain if new alignments were added.

We compared the number of aligning bases in the chains before and after applying RepeatFiller. To this end, we used the coordinates of aligning chain blocks to determine how many bases of the human hg38 assembly align (via at least one chain) to the query species. We used the RepeatMasker repeat annotation for hg38, available at the UCSC Genome Browser [38], to determine how many of the newly-added alignments overlap repetitive elements.

### Generating multiple alignments

Before building multiple alignment, we filtered out low scoring chains and nets requiring a minimum score of 100000. We used Multiz-tba [27] with default parameters to generate two reference-based multiple alignments using the pairwise alignment nets produced with and without RepeatFiller, respectively.

### Conservation analysis

To identify constrained elements, one needs a tree with branch lengths representing the number of substitutions per neutral site. We used four-fold degenerated codon sites based on the human ENSEMBL gene annotation to estimate the neutral branch lengths with PhyloFit [28]. To identify conserved regions, we used PhastCons [28] with the following parameters: rho=0.31; expected-length=45; target-coverage=0.3. To obtain conserved non-exonic regions, we first obtained exonic regions from the human Ensembl and RefSeq annotation (UCSC tables ensGene and refGene). As done before [25], we merged all exonic regions and added 50 bp flanks to exclude splice site proximal regions that often harbor conserved splicing regulatory elements. To obtain Conserved Non-exonic Elements (CNEs), we subtracted these exonic bases and their flanks from all conserved regions.

To compare constraint in genomic regions classified as constraint in only one alignment, we used GERP++ [37] with default parameters (acceptable false positive rate = 0.05) to estimate constraint per genomic position. We denote genomic regions as ‘gained’ if they were classified as constrained by PhastCons only in the multiple alignment generated with RepeatFiller. We denote genomic regions as ‘lost’ if they were classified as constrained only in alignment generated without RepeatFiller (Supplementary Figure 1A). Gained and lost regions were identified using ‘bedtools intersect’ [39]. For each position in ‘gained’ and ‘lost’ non-exonic regions, we computed the RS score (number of rejected substitutions) with GERP++ [37] and calculated the difference between the RS score obtained for the alignment with and without RepeatFiller (Supplementary Figure 1B). These differences are plotted in Figure 5C. Positive differences indicate that the sequences added by RepeatFiller evolve slower than under neutrality, thus increasing the number of rejected substitutions. Differences close to zero indicate that the newly-added sequences evolve as expected under neutral evolution and negative differences indicate that they evolve faster than expected under neutral evolution.

## Data Availability

The multiple genome alignments generated with and without applying RepeatFiller and the respective PhastCons conserved elements are available at https://bds.mpi-cbg.de/hillerlab/RepeatFiller/. The CNEs that differ between both alignments are available in Supplementary Tables 2 and 3. The RepeatFiller source code is available at https://github.com/hillerlab/GenomeAlignmentTools.

## Acknowledgment

We thank the genomics community for sequencing and assembling the genomes and the UCSC genome browser group for providing software and genome annotations. We also thank the Computer Service Facilities of the MPI-CBG and MPI-PKS for their support.

