## Supplement Figure 1 for "RepeatFiller newly identifies megabases of aligning repetitive sequences and improves annotations of conserved non-exonic elements"

This PDF file contains:

- Supplementary Figure 1

Supplementary Tables 1 – 3 are provided as sheets in a separate Excel file.

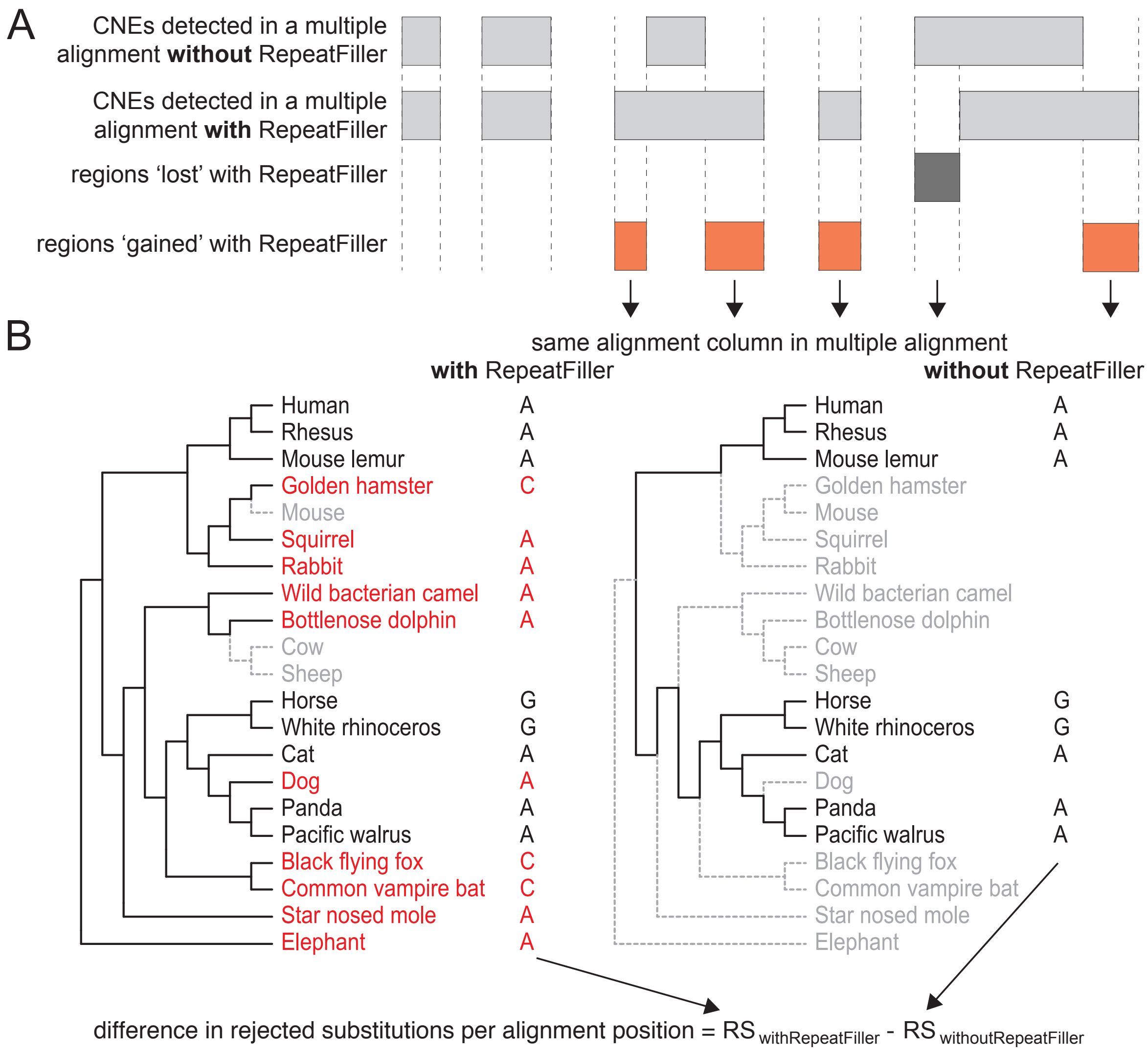


**Supplementary Figure 1:** Comparing constraint in conserved non-exonic regions that were only classified as conserved in the alignment with or without RepeatFiller.

(A) Conserved Non-exonic Elements (CNEs) obtained by PhastCons for alignments with and without RepeatFiller (represented by light grey boxes) are largely identical. However, some of the regions are annotated as conserved either only in the RepeatFiller-subjected alignment ('gained' regions – orange boxes) or only in the alignment without RepeatFiller ('lost' regions – dark grey boxes).

(B) For each position in these variable CNEs, we calculate the number of rejected substitutions (RS) with GERP++, separately for the alignments with and without RepeatFiller. The illustration shows that RepeatFiller adds more aligning sequences (red font). GERP++ computes the number of substitutions expected under neutrality from a phylogenetic tree that is pruned to the aligning species. That means that branches leading to non-aligning species (dashed grey lines) are ignored when computing the number of expected neutral substitutions. The difference in rejected substitutions per alignment column (plotted in Figure 5C) is calculated as the difference of the two RS scores.
